# Comparing the metabolic fates of BALB/c mice maintained on cafeteria-style diets with differential nutritive values

**DOI:** 10.1101/391821

**Authors:** Muhammad Zaid, Fatima Ameer, Ayesha Ali, Zunaira Shoukat, Rida Rashid, Ibrar Iqbal, Nousheen Zaidi

## Abstract

Cafeteria (CAF) diet-fed rodents are shown to provide a robust model of metabolic syndrome and human obesity. The carbohydrate/fat-rich food-items provided to the CAF-diet-model more closely approximate the ultra-processed human diet. However, most of the previous studies applied the commercially available rodent chow-diet for the comparative analyses and labeled it as a healthy-diet. The presented work aims to extend the knowledge on CAF-diet model by exposing the mice to human foods with different nutritional values. Our major goal was to study the metabolic fates of mice maintained on human food-items, which depending upon on their macronutrient compositions are categorized as healthy or unhealthy. BALB/c mice were randomly allocated to one of the three dietary intervention groups, standard chow diet; high-sugar/high-fat-cafeteria (HSHF-CAF) diet; or low-sugar/low-fat-cafeteria (LSLF-CAF) diet, for 5 weeks. The differences in multiple metabolic parameters (including food-/energy /macronutrient-intake, body-weight gain rate, organ-to-body weight ratios, plasma lipid profiles, adipocyte physiology, lipid deposition in metabolic tissues and ectopic fat storage in heart and kidney) were compared among the three intervention groups. We did not observe hyperphagia in mice maintained on CAF-diets. Nonetheless, the CAF-diet-fed mice displayed increased weight-gain-rate, adiposity, and adipocyte hypertrophy when compared to the chow-fed mice. However, the mice maintained on the two cafeteria-style diets displayed similar metabolic profiles, with HSHF-CAF-group displaying slightly higher weight-gain-rate. The HSHF-CAF-and LSLF-CAF-diet induced comparable adiposity in BALB/c mice. Further studies, with longer dietary intervention periods, are required to elucidate the effects of differential CAF-diets on the metabolic health of mice.

## Introduction

Numerous recently published research works –focused at understanding the role of diet in inducing obesity and metabolic syndrome– have utilized rodent models of diet-induced obesity (DIO) through the administration of cafeteria (CAF) diets [1-9]. The CAF-diet consists of highly palatable, energy dense and unhealthy human junk food-items –with high-salt, high-sugar, high-fat and low-fiber content. It has been shown that CAF-diet promotes voluntary hyperphagia that causes rapid weight-gain, dyslipidemia, hyperinsulinemia, hyperglycemia, and glucose intolerance in the rodent models [1, 4, 5, 10-13]. Exposure to the cafeteria-diet was also shown to induce behavioral changes –including reduced activity and low anxiety-like behavior– in rats [5]. Interestingly, it has also been shown that the withdrawal of CAF-diet results in partial recovery from some of the aforementioned metabolic and behavioral alterations associated with CAF-diet-intake [5, 8].

These studies have argued that the CAF-diet-fed rodents can be preferred over the conventional rodent models of high fat (HF)-DIO, that attempt to recapitulate human obesity and metabolic syndrome through the excessive administration of a single macronutrient (i.e. fat) [1, 5, 8]. These HF-regimens do not reflect the nutritional and sensorial diversity of modern human diets. On the other hand, the grocery store-purchased food items provided to the CAF-diet-model more closely approximate the ultra-processed human diet than the commercially available rodent diets [1, 5, 8]. It has been shown that the CAF-diet-fed mice develop metabolic syndrome more severely than the mice maintained on HF-diet [1], creating a phenotype of exaggerated obesity with glucose intolerance and inflammation [1]. The CAF-diet-fed rodents are claimed to provide a more relevant and robust model of human obesity and metabolic syndrome than the rodents consuming monotonous HF-diet [1, 7].

Nonetheless, most of the studies based on the CAF-diet model have utilized the standard chow-fed mice as a control group for the comparative analyses [1, 8]. Even the studies aimed to assess the reversibility of CAF-diet induced metabolic syndrome, by following a “healthy lifestyle” change, labeled the chow-diet as a healthy-diet [8]. We hypothesize that an appropriate control against the CAF-DIO rodent model should also be exposed to the healthier dietary options –with low-sugar and fat content– that are available to human beings. In that way, the impact of modern human diets will be better assessed as both the study and control groups will be consuming ultra-processed food-items with different nutritional values.

In the present study we compared the effects of consuming regular high–sugar/high-fat cafeteria (HSHF-CAF) diet versus “healthier” low–sugar/low-fat cafeteria (LSLF-CAF) diet in mice. The major goal of the study was to compare the metabolic fates of the BALB/c mice maintained on HSHF-CAF-, LSLF-CAF-, or standard-chow-diet. The food-, energy-, and macronutrient-intake were compared among the three diet groups. We also studied various other metabolic parameters –including body-weight-gain-rate, organ-to-body weight ratios, plasma lipid profiles, adipocyte physiology, lipid deposition in metabolic tissues (liver and adipose tissue) and ectopic fat storage in heart/kidney– that have been previously reported to be affected in CAF/HF-diet-fed mice in comparison to the mice maintained on standard-chow diet [4, 14, 15]. Cafeteria diet-fed mice/rats are being increasingly studied and used by the researchers interested in determining the role of aberrant homeostasis in human obesity. The present study significantly contributes to the current understanding of this pertinent model of human obesity and metabolic syndrome.

## Materials and Methods

### Animal experiments and ethics

Age (5-6 weeks) and body weight-matched BALB/c (n = 30, male = 15, female =15) mice were randomly allocated to one of the three dietary intervention groups: standard chow-diet, high-sugar/high-fat cafeteria (HSHF-CAF) diet or low-sugar/low-fat cafeteria (LSLF-CAF) diet (**Figure 1**). The duration of the dietary-intervention period was 5 weeks. The specific food-items chosen for two types of the CAF-diets are listed in **Table 1**. The items that were selected for LSLF-CAF diet had significantly lower carbohydrate, sugar and fat content in comparison to HSHF-CAF diet (**Table 1**). The three experimental diets had varying macronutrient compositions but similar energy densities. Throughout the experiment the mice were allowed *ad libitum* access to water and diet. Food intake was measured twice a week by subtracting the amount of the food left and the initial amount of the food supplied (with spillage taken into account). Energy- and macronutrient-intake from the food consumed was calculated using the known energy and macronutrient content of each food-item (**Table 1**). Body-weights and urinary glucose levels were also recorded twice a week. At the end of the 5-week intervention period, the mice were fasted overnight. The following morning, they were anesthetized with chloroform and after 1-2 minutes blood was drawn from the inferior *vena cava*. The mice were then euthanized by exsanguination. The intra-abdominal white adipose tissue (WAT) (epididymal, mesenteric, retroperitoneal, and perirenal fat pads) was removed and weighed. The liver, kidney, and heart were also removed, weighed, and stored in formalin for histological analyses.

**Figure 1:**
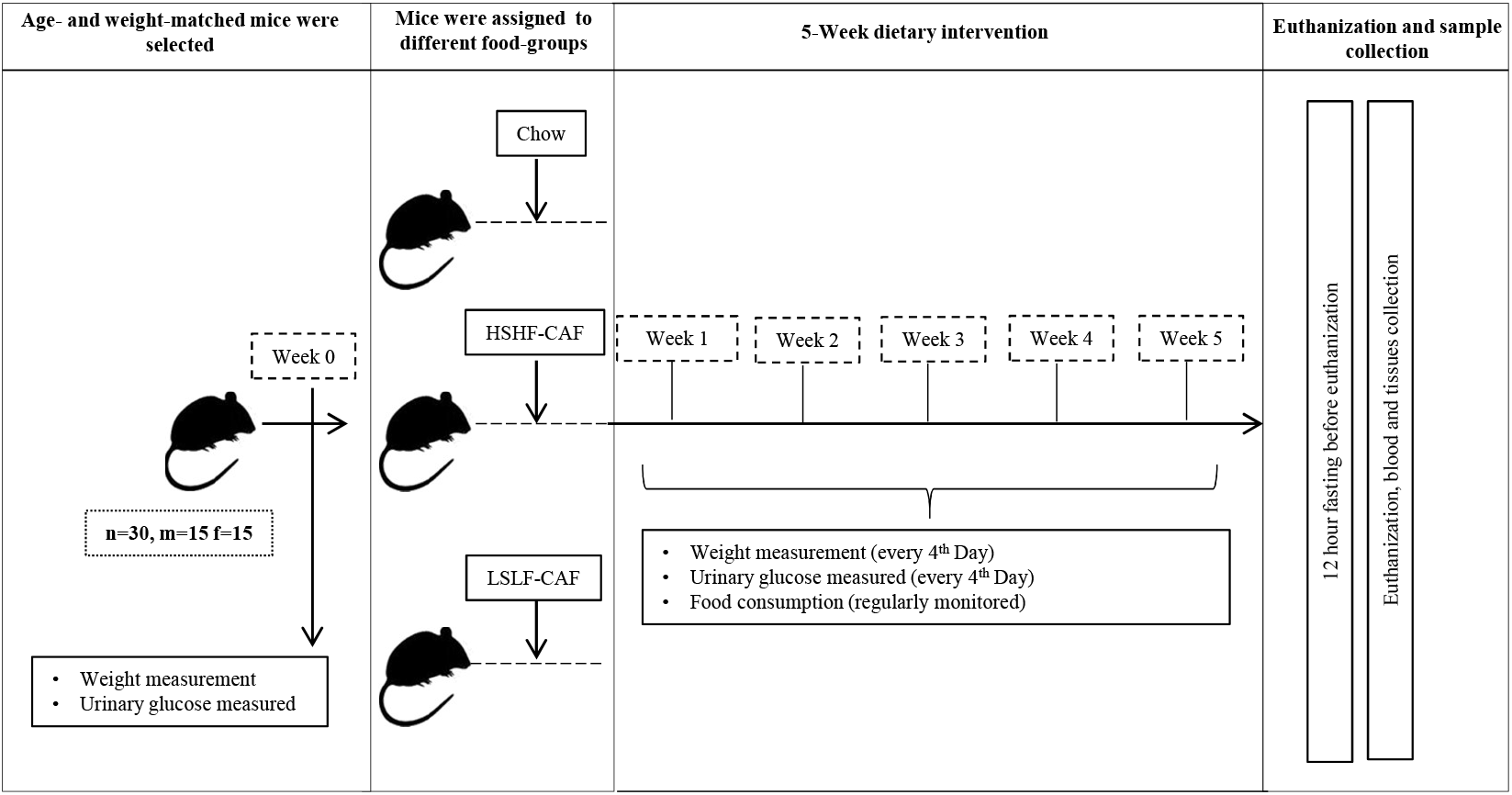
Study Design. Figure summarizes the experimental set-up and data collection points. During a 5-week dietary intervention period, BALB/c mice were studied in three groups: high-sugar/high-fat cafeteria (HSHF-CAF) diet, low-sugar/low–fat cafeteria (LSLF-CAF) diet, or standard-chow diet. All groups included equal number of animals. **Symbols:** n, m and f denote total number of mice, total number of male mice, and total number of female mice respectively.

**Table 1:** Nutritional composition of the food-items provided to the three dietary-intervention groups; high-fat/high-sugar-cafeteria diet, low-fat/low-sugar-cafeteria diet and the standard-chow diet.

| Items | HSHF-CAF-Diet |  |  |  |  | LSLF-CAF-Diet |  |  |  | Control Diet |
| --- | --- | --- | --- | --- | --- | --- | --- | --- | --- | --- |
|  | Cookies/10g | Corn Flakes/10g | Lays chips/10g | White Bread/10g | Chicken nuggets/10g | Sugar free cookies/10g | Bran Flakes/10g | Bran Bread/10g | Red Beans/10g | Chow/10g |
| Energy (kcal) | 47.2 | 36 | 57 | 26.6 | 24.118 | 47.2 | 35.9 | 24.8 | 33.3 | 30 |
| Carbohydrates (g) | 5.76 | 7.94 | 5.3 | 5.1 | 1.7 | 5.66 | 7.3 | 1.7 | 5.9 | 4.44 |
| Sugars (g) | 2.44 | 0.8 | 1 | 0.4 | 0 | 0 | 0.87 | 0.3 | 0.22 | 0.36 |
| Fat (g) | 2.58 | 0.175 | 3.5 | 0.3 | 1 | 3.33 | 0.241 | 0.1 | 0.08 | 0.5 |
| Proteins (g) | 0.54 | 0.665 | 0.714 | 0.8 | 2 | 1.2 | 1.134 | 0.3 | 2.3 | 2.1 |
| Fibers (g) | 0.28 | 0.358 | 0.35 | 0.2 | 0.056 | 0.07 | 1.28 | 0.1 | 1.3 | 0.45 |

Abbreviations: kcal, kilocalories; HSHF-CAF, high-sugar/high-fat cafeteria; LSLF-CAF, low-sugar/low-fat cafeteria.

The study protocols were approved by the Biosafety and Bio-resource Committee, University of the Punjab. All efforts were made that the use of animals causes the least possible amount of suffering.

### Histological Analyses

The organs/tissues (adipose tissue, liver, heart and kidney) were removed and fixed by immersion in 4 % paraformaldehyde, dehydrated, and embedded in paraffin. Thin sections of 3–5 μm were cut and mounted on silanized slides. The slides were stained with hematoxylin and eosin according to the standard routine laboratory procedures. For histopathological analyses, slides were examined by light microscopy and photomicrographs were taken using an HD1500T Meiji American Technology Camera mounted on an Euromex microscope.

### Determination of Adipocyte Size

The same (reteroperitoneal) region of the fat pad was used for all animals to minimize cell size variation due to differences in anatomical location. Mature white adipocytes were identified by their characteristic multilocular appearance. Total areas of adipocytes were traced and analyzed with Adiposoft (version 1.13) plugin within ImageJ software (MATLAB). White adipocyte areas were measured in 50-60 or more cells per mouse in the respective groups.

### Determination of plasma lipid levels

Plasma total cholesterol levels were spectrophotometrically determined using commercially available kit (Analyticon Biotechnologies AG, 4046). Plasma triglyceride levels were determined using commercially available kit for quantitative determination of triglycerides in human serum/plasma (Analyticon Biotechnologies AG, Catalogue # 5052).

### Measurements of lipids in tissues

Lipids were extracted from organ slices using a methanol/chloroform extraction method as previously described [16]. Total cholesterol content in the lipid extracts was spectrophotometrically determined using commercially available kit (Analyticon Biotechnologies AG, 4046) against a calibration-curve generated using known concentrations of cholesterol standard (SUPELCO, 47127-U). Total triglyceride content in the lipid extracts was spectrophotometrically determined using commercially available kit (Analyticon Biotechnologies AG, Catalogue # 5052) against a calibration-curve generated using known concentrations of triglyceride standard (SUPELCO, 17811-1AMP).

### Statistical analysis

The differences between groups were analyzed by repeated measures ANOVA or t-test (paired or unpaired), where applicable. Results were analyzed by GraphPad Prism Software, Version VI. P-values <0.05 were considered statistically significant and indicated when different.

## Results

### Effects of different diets on feeding-patterns in BALB/c mice

Mice were fed (*ad libitum*) with standard-chow-, high-sugar/high-fat-cafeteria (HSHF-CAF)-, or low-sugar/low-fat-cafeteria (LSLF-CAF)-diet. Food- and energy-intake were regularly monitored during the intervention-period (5 weeks). Periodic fluctuations in food- and energy-intake were noted in mice from all the three diet-groups (**Figure 2a-f**). However, no significant difference was observed between the mice on standard-chow-, HSHF-CAF-, or LSLF-CAF-diet (**Figure 2a-f**). We also compared the intake of macronutrients among the three diet groups. It was observed that the mice on HSHF-CAF-diet persistently consumed more carbohydrates and sugars than the mice on standard-chow- or LSLF-CAF-diet (**Figure 3a-b**), whereas the carbohydrates/sugar-intake was not significantly different between standard-chow- and LSLF-CAF-group. The fat-intake was significantly different among the three groups –with HSHF-CAF-group consuming highest and standard-chow-group consuming the lowest fat content (**Figure 3c**). Protein-intake was significantly higher in standard-chow-group when compared to LSLF-CAF- or HSHF-CAF-group (**Figure 3d**), whereas the protein-intake was not significantly different between HSHF-CAF- and LSLF-CAF-group. In order to match the protein content of the CAF-diets utilized in the previous studies, HSHF-CAF- and LSLF-CAF-regimens respectively included processed meat and boiled red beans. However, the animals did not consume these food-items.

**Figure 2:**
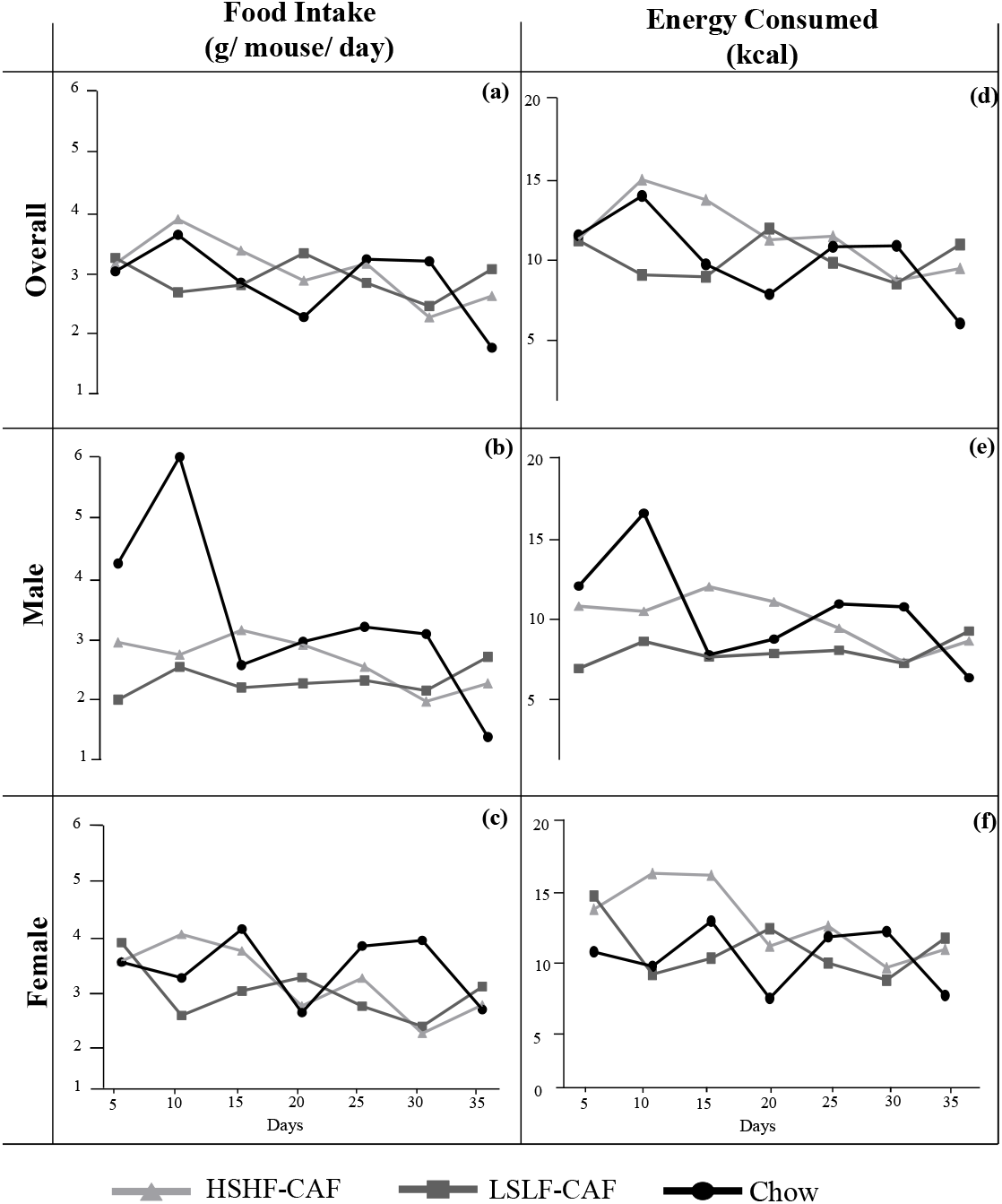
Food- and energy-intake in mice maintained on standard-chow-diet or cafeteria-diets with differential nutritive values. Line-graphs represent food-intake (g/mouse/day) in (a) overall (b) male (c) female mice and energy-intake (kcal) in (d) overall (e) male (f) female mice maintained on standard-chow-, HSHF-CAF-, or LSLF-CAF-diet. Food- and energy-intake were regularly monitored during the intervention period (5 weeks). Data were analyzed using repeated measures ANOVA (n = 10, males = 5, and females = 5 for each group). **Abbreviations**: kcal, kilocalories; HSHF-CAF, high-sugar/high–fat cafeteria; LSLF-CAF, low-sugar/low-fat cafeteria.

**Figure 3:**
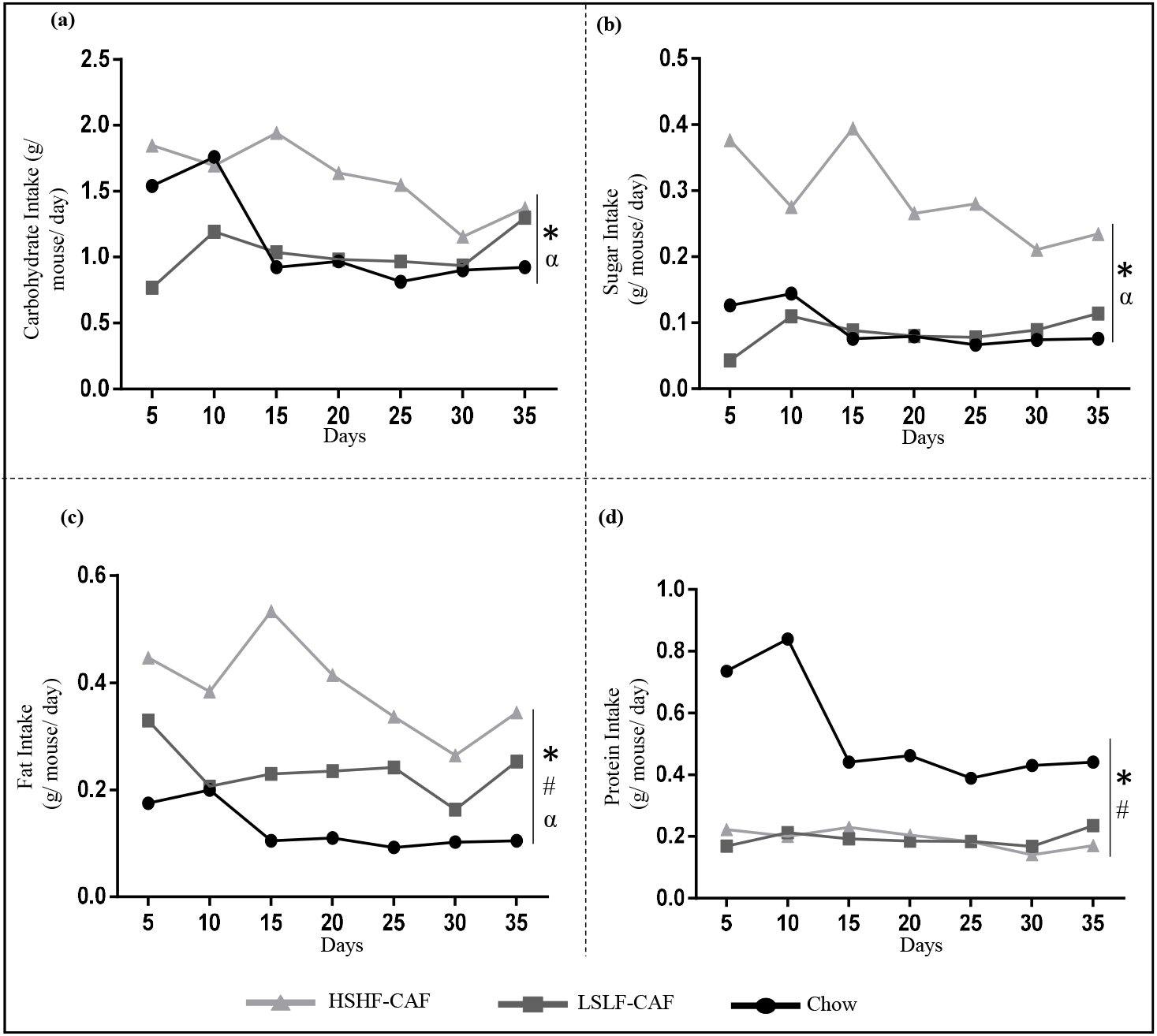
Macronutrient-intake in mice maintained on standard-chow-diet or cafeteria-diets with differential nutritive values. Line-graphs compare. (a) carbohydrate- (b) sugar- (c) fat- and (d) protein-intake in mice maintained on standard-chow-, HSHF-CAF-, or LSLF-CAF-diet. Macronutrients-intakes were accordingly calculated by applying net food-intake (see figure 2) and macronutrient-compositions of diets. Data were analyzed using Student’s t-test. Symbols of statistical significance (P <0.05) *, #, and α respectively refer to the comparison of standard-chow-diet vs. HSHF-CAF-diet, standard-chow-diet vs. LSLF-CAF-diet, and HSHF-CAF-diet vs. LSLF-CAF-diet **Abbreviations:** HSHF-CAF, high-sugar/high–fat cafeteria; LSLF-CAF, low-sugar/low-fat cafeteria.

### Metabolic consequences of consuming standard-chow-, HSHF-CAF-, or LSLF-CAF-diet

We monitored the body-weights of mice maintained on standard-chow-, HSHF-CAF- or LSLF-CAF-diet during the intervention-period of 5 weeks. The mice on standard-chow-diet gained weight at a significantly slower rate than the mice on any of the two CAF-diets. The mice maintained on the two cafeteria-style diets displayed no significant difference -with HSHF-CAF displaying slightly higher weight-gain-rate (**Figure 4a**).

**Figure 4:**
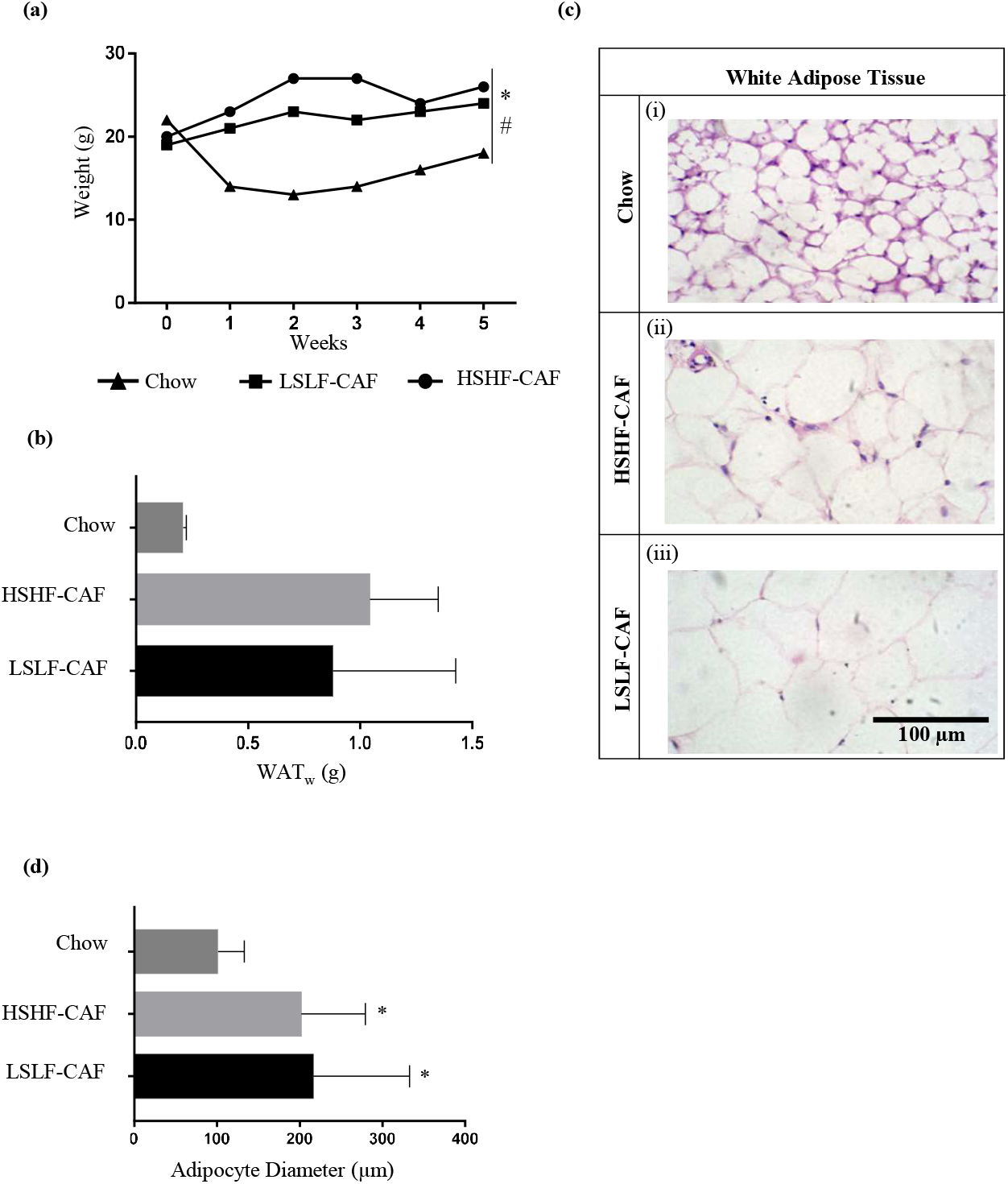
Body weight gain, increased adipose tissue mass, and adipocyte hypertrophy in mice on cafeteria-diets with differential nutritive values. **(a)** Line-graphs represent body-weights (grams) of mice maintained on standard-chow-, HSHF-CAF-, or LSLF-CAF-diet. Body-weight of each animal was recorded twice a week during the intervention period (5 weeks). Data were analyzed using Student’s t-test (n = 10 for each group). Symbols of statistical significance (P <0.05) *, #, and α respectively refer to the comparison of standard-chow diet vs. HSHF-CAF-diet, standard-chow-diet vs. LSLF-CAF-diet and HSHF-CAF-diet vs. LSLF-CAF-diet. **(b)** Total weight of white adipose tissue (WAT_W_) (epididymal, mesenteric, retroperitoneal, and perirenal fat pads) from mice maintained on standard-chow-, HSHF-CAF-, or LSLF-CAF-diet.**(c)** Histological analysis of retroperitoneal white adipose tissue from mice maintained on standard-chow-, HSHF-CAF-, or LSLF-CAF-diet. Bars indicate 100 μm. **(d)** Cell size of retroperitoneal adipocytes from mice maintained on standard-chow-, HSHF-CAF-, or LSLF-CAF-diet. Values are expressed as the mean ± SD (n = 30-40). Data were analyzed using Student’s t-test. *Significant difference comparing standard-chow-diet with HSHF-CAF-diet or LSLF-CAF-diet (P <0.05). **Abbreviations:** HSHF-CAF, high-sugar/high-fat cafeteria; LSLF-CAF, low-sugar/low-fat cafeteria.

Total white adipose tissue mass (WAT_W_) was significantly lower in mice on standard-chow-diet when compared to the mice on HSHF-CAF- and LSLF-CAF-diet (**Figure 4b**). However, no significant difference was observed between the HSHF-CAF and LSLF-CAF groups (**Figure 4b**). Histological analyses of reteroperitoneal fat pads (**Figure 4c**) and quantitation of adipocyte size (**Figure 4d**) revealed that adipocytes from HSHF-CAF- and LSLF-CAF-group were significantly larger than those from chow-group. No significant difference was observed between the HSHF-CAF- and LSLF-CAF-group (**Figure 4c-d**). In contrast to the previous studies, we did not observe any significant difference between plasma lipid profiles of mice in any of the three diet groups (**Table 2**).

**Table 2:** Plasma lipid profiles of the mice maintained on standard-chow-, HSHF-CAF-, and LSLF-CAF-diets.

| Parameters | Male |  |  | Female |  |  |
| --- | --- | --- | --- | --- | --- | --- |
|  | **Chow-Fed**<br>**Mice**<br>**(n=5)** | **HSHF-CAF-Fed**<br>**Mice**<br>**(n=5)** | **LSLF-CAF-Fed**<br>**Mice**<br>**(n=5)** | **Chow-Fed**<br>**Mice**<br>**(n=5)** | **HSHF-CAF-Fed**<br>**Mice**<br>**(n=5)** | **LSLF-CAF-Fed**<br>**Mice**<br>**(n=5)** |
| Total Cholesterol (mg/dL) | 68.74 ± 31.47 | 56.31 ± 24.75 | 76.75 ± 25.01 | 76 ± 27.055 | 67.35 ± 6.99 | 42.18 ± 16.88 |
| Triglycerides (mg/dL) | 94.24 ± 49.5 | 62.22 ± 20.90 | 155.16 ± 121.74 | 94 ± 39.15 | 125.82 ± 42.71 | 54.83 ± 15.97 |

*Values are presented as mean ± SD. Significant difference between two groups was assessed by Student's t-test (\*P <0.05). Abbreviations: HSHF-CAF, high-sugar/high-fat cafeteria; LSLF-CAF, low-sugar/low-fat cafeteria.*

### Impact of standard-chow-, HSHF-CAF-, or LSLF-CAF-diet on organ-weights, -pathology and -lipid deposition

In accordance with a previous study [4] we did not observe any significant difference in organ to body-weight ratios among the mice on standard-chow-, HSHF-CAF-, or LSLF-CAF-diet (**Table 3**). It has been previously reported that cafeteria-style diet induces organ damage in mice. Here, we did not observe any significant differences in pathology of liver, heart, or kidney from mice on standard-chow-, HSHF-CAF-, or LSLF-CAF-diet (**Figure 5**). We also examined lipid-deposition in liver and adipose tissues of mice from the three diet groups. As shown in Table 4 no significant differences were observed in cholesterol and triglyceride deposits within these tissues. The storage of ectopic fat in the key target-organs of cardiovascular control –kidney and heart– was also compared between the three diet groups. Again, no significant differences were observed among the mice maintained on standard-chow-, HSHF-CAF-or LSLF-CAF-diet (**Table 5**).

**Table 3:** Organ-to-body weight ratios of the mice maintained on standard-chow-, HSHF-CAF- and LSLF-CAF-diet.

| Parameters | Male |  |  | Female |  |  |
| --- | --- | --- | --- | --- | --- | --- |
|  | **Chow-Fed Mice (n=5)** | **HSHF-CAF-Fed Mice (n=5)** | **LSLF-CAF-Fed Mice (n=5)** | **Chow-Fed Mice (n=5)** | **HSHF-CAF-Fed Mice (n=5)** | **LSLF-CAF-Fed Mice (n=5)** |
| Liver-to-Body Weight | 0.046 ± 0.006 | 0.046 ± 0.008 | 0.034 ± 0.011 | 0.047 ± 0.004 | 0.024 ± 0.017* | 0.043 ± 0.007 |
| Kidney-to-Body Weight | 0.015 ± 0.001 | 0.018 ± 0.006 | 0.018 ± 0.0025 | 0.014 ± 0.002 | 0.012 ± 0.001 | 0.013 ± 0.003 |
| Heart-to-Body Weight | 0.005 ± 0.000 | 0.0056 ± 0.000 | 0.0053 ± 0.001 | 0.004 ± 0.001 | 0.004 ± 0.001 | 0.004 ± 0.001 |

*Values are presented as mean ± SD. Significant difference between two groups was assessed by Student's t-test. Statistical significance (\*P <0.05) is indicated when different.*
Abbreviations: HSHF-CAF, high-sugar/high-fat cafeteria; LSLF-CAF, low-sugar/low-fat cafeteria.

**Figure 5:**
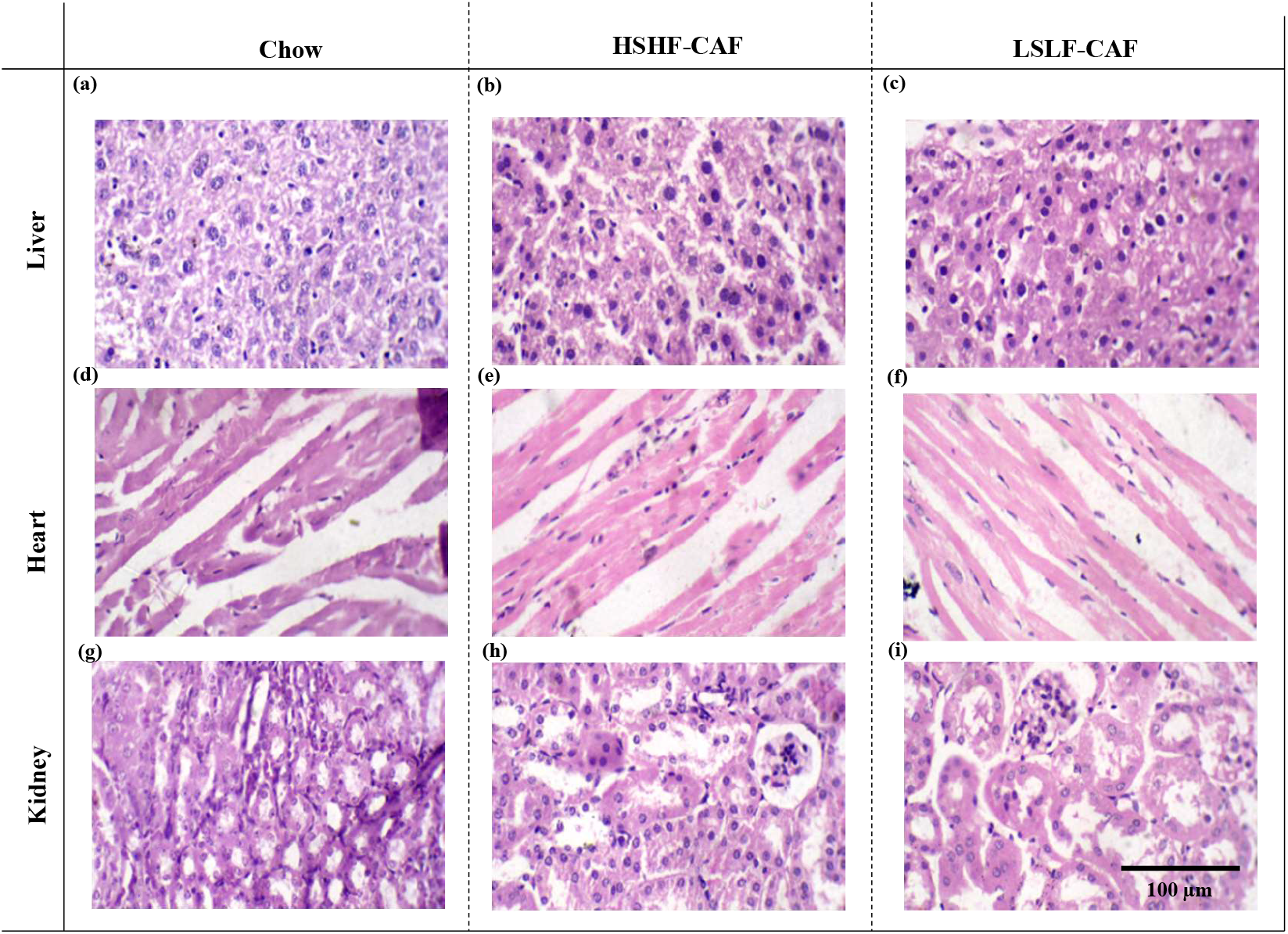
Organ (liver, heart and kidney) pathology in mice maintained on standard-chow-diet or cafeteria-diets with differential nutritive values. Histological analyses of **(a-c)** liver **(d-f)** heart and **(g-i)** kidney from mice respectively maintained on standard-chow diet, HSHF-CAF-, or LSLF-CAF-diet. Organ sections were stained with hematoxylin and eosin **(H&E)**. Bars indicate 100 μm. **Abbreviations:** HSHF-CAF, high-sugar/high-fat cafeteria; LSLF-CAF, low-sugar/low-fat cafeteria.

**Table 4:**
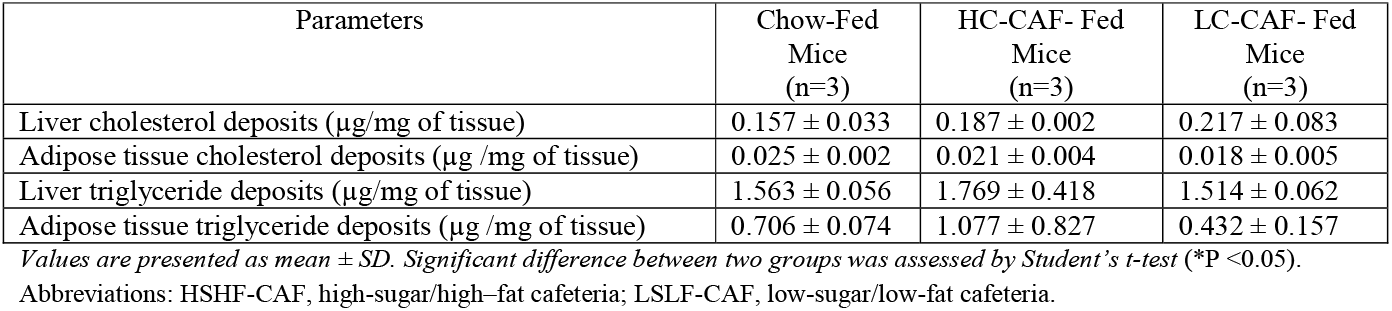
Impact of various cafeteria-style diets on lipid deposition in metabolic tissues.

**Table 5:** Impact of various Cafeteria-style diets on ectopic fat storage in the key target-organs of cardiovascular control (heart and kidney).

| Parameters | Chow-Fed<br>Mice<br>(n=3) | HC-CAF- Fed<br>Mice<br>(n=3) | LC-CAF- Fed<br>Mice<br>(n=3) |
| --- | --- | --- | --- |
| Kidney cholesterol deposits ($\mu\text{g}$ /mg of tissue) | $0.388 \pm 0.059$ | $0.333 \pm 0.080$ | $0.372 \pm 0.045$ |
| Heart cholesterol deposits ($\mu\text{g}$ /mg of tissue) | $0.146 \pm 0.016$ | $0.237 \pm 0.025$ | $0.138 \pm 0.015$ |
| Kidney triglyceride deposits ($\mu\text{g}$ /mg of tissue) | $1.620 \pm 0.211$ | $1.468 \pm 0.099$ | $1.676 \pm 0.264$ |
| Heart triglyceride deposits ($\mu\text{g}$ /mg of tissue) | $1.103 \pm 0.346$ | $0.994 \pm 0.137$ | $0.967 \pm 0.553$ |

*Values are presented as mean ± SD. Significant difference between two groups was assessed by Student's t-test (\*P <0.05).*
Abbreviations: HSHF-CAF, high-sugar/high-fat cafeteria; LSLF-CAF, low-sugar/low-fat cafeteria.

## Discussion

In the recent years the cafeteria diet-fed rats/mice have emerged as pertinent animal models for studying human obesity and metabolic disorders [1-9]. To the best of our knowledge, none of the previous studies have compared the effects of regular CAF-diet – containing high-fat and high-sugar content-with the “healthier” grocery-purchased human food-items that have lower fat and sugar content. The presented study compares the metabolic fates of mice maintained on standard-chow-, high-sugar/high-fat-cafeteria (HSHF-CAF)-, or low-sugar/low-fat-cafeteria (LSLF-CAF)-diet.

We observed that the food-intake did not significantly vary among the three diet-groups. This observation contradicts with the previous studies that have shown that the mice/rats maintained on CAF-diets develop voluntary hyperphagia and display significantly higher food-intake [1, 2, 4]. Some of the previous works have presented contradictory data on the feeding patterns in CAF-fed rats [2, 17]. It has been argued that the differences in the feeding patterns could be attributed to the variety of food-items provided to the CAF-diet groups. One of the studies that provided wider range of food-items showed that the CAF-fed rats ate larger meals than chow-fed rats across the entire duration of the study [2]. Another study, in which the rats were exposed to lesser number of cafeteria-style food-items, displayed that the differences in the food-intake patterns between the CAF- and chow-diet groups declined following the initial phase of elevated-intake [17]. The lack of hyperphagia in our CAF-diet groups might also be due to the lesser number of food-items that were utilized for this study as the variety in diet is shown to reduce sensory-specific satiety thereby enhancing the food intake [17, 18].

Despite the fact that the animals consumed similar amounts of food the distinctive macronutrients composition of the experimental diets resulted in differential macronutrient-intake in the three diet-groups throughout the course of the study. These differences resulted in increased weight-gain, adiposity and adipocyte hypertrophy in the mice on CAF-diets when compared to the chow-fed mice. However, despite the obvious difference in the nutritive values of HSHF-CAF and LSLF-CAF diets no significant metabolic differences were observed between the mice consuming these diets. Additionally, in contrast to the previous works we did not observe any significant differences in organ-to-body weight ratios, plasma lipid profiles, lipid deposition in metabolic tissues (liver and adipose tissue), and ectopic fat storage in heart and kidney, among the three diet groups [14]. Moreover, no significant differences in organ pathology were noted in the mice maintained of CAF-diets as suggested by the previous works [7].

The contradictions between the previous studies and our data might be attributed to multiple factors. Most importantly, the specific food-items that were selected for our study were different from the previous works. Moreover, as discussed above the food items included in our CAF-diet regimens provided a relatively narrow range of dietary options to the animals. These factors might have contributed to the lack of hyperphagia in our CAF-diet groups, hence preventing the emergence of severe forms of metabolic abnormalities that were observed in the previous works [1, 2, 4]. The other factor that might have contributed to these inconsistencies is the shorter dietary intervention period. Most of the previous works exposed the animals to different types of diets for longer intervention periods (15-30 weeks) [1, 2, 19].

Our observations suggest that the application of CAF-diet model warrants careful considerations regarding the choice, variety and exposure time of the CAF-diets. Some of the earlier works have also raised doubts regarding the aptness of the CAF-diet-fed mice as models of human obesity on the similar grounds. It was argued that the nutritive and nonnutritive components of the food-items are not well defined in CAF-diets. In addition, the animal may choose a different selection of foods each day that may cause differential metabolic consequences in different studies or animals within the same study group [20]. Hence, it can be suggested that to make an appropriate CAF-DIO model and generate reproducible data the selection of food-items should be standardized.

The mice are widely used as a model of genetic obesity [21-23]. However, currently the studies focused at understanding the role of diet in inducing obesity predominantly use rats as model organisms. One of the major reasons of this occurrence is that the mouse strains show different biological responses to stress and chronic illness [24-27]. For instance, both C57BL/6J and BALB/c mice maintained on high-fat diet display marked increase in body-weights but dissimilarities in various clinical and pathological parameters [28]. In the recent past, several studies focused on CAF-diet induced obesity have utilized mice as a model organism. The extraction of relevant information from scientific literature, for designing future CAF-model based studies, requires emphasis on rodent- and strain-types utilized in the previous works.

### 2. Conclusions

One of the most remarkable observations of the presented work is that the CAF diets - either with low-sugar/low-fat or high–sugar/high-fat content– caused increased weight gain and adiposity in mice in comparison to the standard-chow-diet. The presented study significantly adds to the current understanding of CAF-diet induced obesity in rodent models. Further rodent-based studies, with longer dietary intervention periods, are required in order to better understand the effects of differential CAF-diets on adiposity, obesity and induction of metabolic syndrome.

## Acknowledgement

This work was supported by the Higher Education Commission of Pakistan (Project 2505/R&D/11-2670) and University of the Punjab (Annual Research Grant) (Principal Investigator: Nousheen Zaidi).

---

**Supplementary Table 1:**
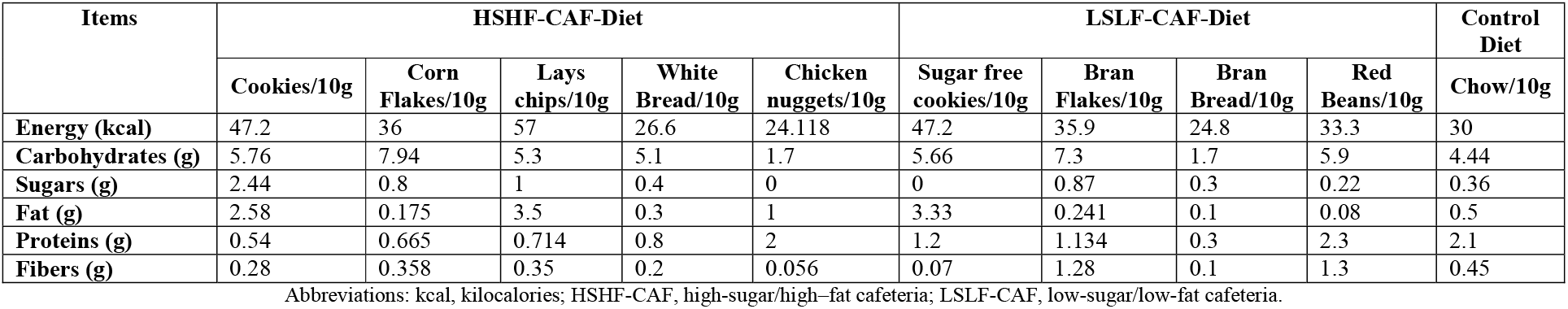
Nutritional composition of the food-items provided to the three dietary-intervention groups; high-fat/high-sugar-cafeteria diet, low-fat/low-sugar-cafeteria diet and the standard-chow diet.

## Abbreviations

HSHF-CAF: High-sugar/high-fat-cafeteria
LSLF-CAF: Low-sugar/low-fat-cafeteria
CAF: Cafeteria
HF: High-fat
DIO: Diet induced obesity.

